# Development of an optimized, non-stem cell line for intranasal delivery of therapeutic cargo to the central nervous system

**DOI:** 10.1101/2023.08.16.553513

**Authors:** Ali El-Ayoubi, Arsen Arakelyan, Moritz Klawitter, Luisa Merk, Siras Hakobyan, Irene Gonzalez-Menendez, Leticia Quintanilla-Fend, Per Sonne Holm, Wolfgang Mikulits, Matthias Schwab, Lusine Danielyan, Ulrike Naumann

## Abstract

Neural stem cells (NSC) are considered to be valuable candidates for delivering a variety of anti-cancer agents to brain tumors, including oncolytic viruses. However, owing to the previously reported tumorigenic potential of NSC cell line after intranasal administration (INA), here we identified the human hepatic stellate cell line (LX-2) as a cell type capable of longer resistance to replication of oncolytic adenoviruses (OAV) as therapeutic cargo, and being non-tumorigenic after INA. Our data show that LX-2 cells can longer withstand the OAV XVir-N-31 replication and oncolysis than NSCs. By selecting the highly migratory cell population out of LX-2, an offspring cell line with a higher and more stable capability to migrate was generated. Additionally, as a safety backup, we applied genomic HSV-TK integration into LX-2 leading to the high vulnerability to Ganciclovir. Histopathological analyses confirmed the absence of neoplasia in the respiratory tracts and brains of immuno-compromised mice 3 months after INA of LX-2 cells. Our data suggest that LX-2 is a novel robust and safe cell line for delivering anti-cancer and other therapeutic agents to the brain.

## 1. Introduction

Neurodegenerative diseases such as Alzheimer’s (AD) or Parkinson’s disease (PD), and malignant brain tumors remain amongst the greatest challenges in modern therapy of central nervous system (CNS) disorders. Stem cell-based therapies have been used so far to provide neuroprotection and therefore to improve neurological functions in neurodegenerative diseases (1), as well as to deliver a variety of anti-cancer agents to brain tumors (2). In particular neural (NSCs) or mesenchymal stem cells (MSCs) have been employed as cell candidates to treat brain diseases as they provide several advantages: (i) NSCs produce a wide repertoire of neuroprotective and trophic factors (3) and therefore can help counteract degeneration-caused neuronal damage and cell death (4); (ii) stem cells (SCs) can be easily genetically modified to express a wide variety of genes that are either neuroprotective or can inhibit tumor growth; (iii) stem cells can also be loaded with therapeutic cargo such as nanoparticles or oncolytic viral vectors (5). However, the major disadvantage of using stem cells (SC) is their tumorigenic potential as has been shown in immunocompromised mice (6-8). This limits the clinical translation of SC cell lines and calls for identifying more differentiated cells that can be used as carriers of therapeutic agents to treat brain diseases.

The blood-brain barrier (BBB) is one of the major limiting factors in the delivery of a variety of drugs and biologics (including stem cells) to the CNS via systemic, such as intravenous, administration. On the other hand, the invasiveness of stereotactic or intrathecal injection limits their implementation in the therapy of neurological disorders. Glioblastoma (GBM) are high-grade and very aggressive adult brain tumors with a median survival of patients less than 22 months even at optimal care and treatment (9). The immunosuppressive tumor microenvironment (TME) of GBMs that prevents tumor immune surveillance, its radio- and chemoresistance and infiltrative growth leading to the spreading of GBM cells throughout the brain, as well as tumor cell heterogeneity are the major contributors to the extremely poor survival of patients (for review see (10)).

Numerous encouraging preclinical studies on oncolytic virotherapy (OVT) of GBM have brought forward this strategy to clinical translation with a first in human trial demonstrating the safety of OVT in humans. However, despite high expectations from preclinical research, OVT bears several drawbacks underpinning the scarce outcomes in clinical efficacy trials (11). Firstly, due to the poor BBB permeability to OVs and to the patient’s immune system-induced rapid inactivation of systemically applied viruses, OVs must be administered intratumorally which limits repeated administration in treating tumor recurrency. Secondly, OVs cannot replicate in tumor-surrounding non-neoplastic cells, which restricts the viral spreading to or near to the virus injection site. Such tumor selectivity of OVs allows avoiding neurotoxicity to surrounding healthy tissue. On the other hand, it permits invaded tumor cells to escape the OV-induced killing, especially when they are located remotely from the original tumor or the main locus of recurrence (12). In this sense, using cells as vehicles delivering OVs to the tumor site and a delivery method that will allow targeting invaded tumor cells within the healthy brain tissue will help to address the aforementioned drawbacks of OVT.

Since the first discovery of intranasal cell delivery to the brain (13) a wide array of preclinical data over the last decade has successfully proven intranasal administration (INA) to be non-invasive, targeted and efficacious administration route allowing a big variety of therapeutic agents, such as drugs, plasmids, peptides, viruses, metals, and OV-loaded cells to bypass the BBB and be delivered to the CNS (14-16). Furthermore, in contrast to stereotactic brain injection, INA can be performed repeatedly and if compared to intravenous administration, helps avoid systemic distribution of an applied drug. Previous own research demonstrated targeted intranasal delivery of mesenchymal stem cells (MSC) to the tumor site in a mouse model of GBM (17). A first-in-human trial has recently proven intranasal delivery of MSCs to be safe (18) empowering this administration route to be employed in cell-based therapies of a variety of neurologic disorders using primary autologous or allogenic stem cells. However, when it comes to the implementation of cell lines by non-invasive administration routes such as INA or intravenous administration, an accurate histopathological analysis of the respiratory tract should be performed to exclude the tumorigenic potential of a cell line in case of unintended delivery of a cell portion to the lung.

In this work we sought to establish for OVT a cell line originated from human somatic differentiated cells with high permissiveness to the viral uptake and phenotypical stability over long periods of cultivation. A pertinent option in our opinion was the genetically unmodified, spontaneously immortalized cell line LX-2 that originated from human hepatic stellate cells (HepSCs) (19). HepSCs are liver perisinusoidal cells located in the subendothelial space between the surface of hepatocytes and endothelial cells (20). Due to their remarkable functional plasticity HepSCs contribute to both, fibrogenesis and repair processes in the liver (21). Under stimulation induced by hypoxia or transforming growth factor beta (TGF-β), HepSCs undergo a transition from a quiescent to an activated phenotype and then show elevated proliferation, increased cell motility as well as adhesion and the expression of several cytokines (22-25). Therefore, we suggested that LX-2 cells might be feasible cells that can be used to rapidly transport OVs or potentially other therapeutic agents to the brain by INA. For the proof of principle, we used the OV XVir-N-31 as a therapeutic cargo and aimed to characterize the properties of LX-2 to resist viral replication, to produce infectious virus particles and at the same time to retain their migratory capacities. XVir-N-31 (also named Ad-Delo3-RD) is an oncolytic, genetically modified adenovirus that has been tested in several mouse tumor models including GBM. Intratumoral injection of XVir-N-31 in GBM bearing mice significantly prolonged the survival of animals, however, failed to reach curative intent (26-30).

## 2. Methods

### 2.1. Cells, cell lines and cell culture

LX-2 cells, a kind gift from Scott Friedman (Division of Liver Diseases, The Icahn School of Medicine at Mount Sinai, NY, USA; Cellosaurus ID: CVCL_5792) were cultivated in DMEM containing 2% fetal calf serum (FCS), 1% glutamine and 1% penicillin/streptomycin (P/S, all from Sigma Aldrich, Darmstadt, Germany) and were described in detail in (19). LX-2^mCherry^ (PAR), LX-2^mCherry^ fast running (FR) and LX-2^mCherry^ fast running and HSV-TK expressing (FR/TK) cells were generated by the infections of the cells with LV-mCherry, followed by the infection with LV-TK, and subsequent puromycin selection. M1-4HSC mouse HepSCs (31) were kindly provided by W. Mikulits (Medical University of Vienna, Vienna Austria). HB1.F3 v-myc-immortalized human NSCs (32) were from H.J. Lee (College of Medicine and Medical Research Institute, Chungbuk National University, Republic of Korea; Cellosaurus ID: CVCL_LJ44). Both cell lines were cultured in DMEM containing 10% FCS, 1% glutamine, 1% P/S and 1% non-essential amino acids (MEM-NEAA, Gibco/Thermo Fisher Scientific, Weil am Rhein, Germany). LN-229 and U87MG (Cellosauro ID: CVCL_0393, CVCL_0022) human glioma cells were a kind gift from N. Tribolet (Geneva, Switzerland) and are described in detail in (33). LN-229 GFP expressing cells were produced by infection with Lenti-GFP (Amsbio, Frankfurt/Main, Germany). HEK293FT were from Thermo Fisher Scientific (Weil am Rhein, Germany; Cellosaurus ID: CVCL_6911) and HEK293 cells from Microbix (Mississauga, ON, Canada; Cellosaurus ID: CVCL_0045). HB1.F3 v-myc-immortalized human NSCs, LN-229, U87MG and HEK293 cells were cultured in DMEM containing 10% FCS, 1%P/S. The R49 glioma stem cell (GSC) line was kindly provided by C. Beier (University Odense, Denmark), is described in (34) and was maintained as tumor spheres in stem cell-permissive Dulbecco’s modified Eagle’s medium/F12 medium (Sigma) supplemented with human recombinant epidermal growth factor (EGF; BD Biosciences), human recombinant basic fibroblast growth factor (bFGF; R&D Systems Europe, Ltd.), human leukemia inhibitory factor (Millipore; 20 ng/ml each) and 2% B27 supplement (Thermo Fisher Scientific, Inc.). All cells were cultured at 37°C in a humidified, 5% CO_2_ containing atmosphere. All human cell lines underwent cell line authentification in May, 2023 (Eurofins Genomics Europe Shared Services GmbH, Ebersberg, Germany; Suppl. Fig. 6). For the generation of supernatants, the cells were grown in serum-deprived medium for 48h. For the generation of conditioned medium, supernatants from semi-confluent cells were collected and clarified from cell debris by centrifugation. Conditioned medium was stored at −80°C until usage. To determine the growth rate or the cytotoxicity of the cells after GVC treatment (Sigma Aldrich), the cells were seeded in triplicates in 96 well plates. Cell density was measured by staining the cells with crystal violet as described (35).

### 2.2. Selection of LX-2 “fast running” cells

LX-2 ^“^fast running” cells (FR) were generated by our previously developed and characterized method of selection of highly migratory subpopulation of cells (17). Briefly, 1 × 10^5^ LX-2 cells were seeded in the top chamber of an 8 μm pore migration cassette (Corning, Kaiserslautern, Germany) and were allowed to migrate for 4,5 h. As attraction medium, conditioned medium from LX-2 cells was used. Migrated LX-2 cells on the bottom part of the membrane were collected by trypsination and were allowed to grow. This procedure was repeated four times. Finally, the selected cells were expanded in culture, and their migration capacity was tested over several passages.

### 2.3. Determination of cell motility and invasion

Cell migration was performed as described. Cells were allowed to migrate for 5 h against conditioned medium, or against 200.000 GBM or GSCs seeded the day before on the bottom of the lower chamber. Migrated cells on the bottom side of the membrane were either stained with crystal violet or were collected by trypsination and counted using a cell titer blue staining kit (CTB; Sigma-Aldrich). For measuring invasion, a commercial matrigel invasion assay (R&D Systems, Abington, UK) was used according to the manufacturer’s protocol. Live cell imaging of migrating cells was performed by seeding 50.000 cells in triplicates in 6 cm petri dishes. After attachment single cells (9-12 cells per dish) were tracked every 10 minutes during a period of 24 h using the CytoSmart and Axion Imaging Systems (CytoSMART Technologies, Eindhoven, The Netherland; Axion BioSystems, Atlanta, Georgia, USA). Velocity, accumulated and Euclidian distances were determined by manual tracking using the Image J “Manual Tracking” plug in (http://rsb.info.nih.gov/iJ/plugins/manual-tracking.html; Fiji, (36)). Visualization and data analysis were performed using the Chemotaxis and Migration Tool 2.0 (Ibidi GmbH, Martinsried, Germany).

### 2.4. Lentivirus cloning, preparation and infection

The lentiviral shuttle vector coding for mCherry (LV-mCherry) was a from Addgene (#36084, Cambridge, MA, USA). LV-TK, containing the herpes simplex virus thymidin kinase coding sequence (HSV-TK), was generated by cloning HSV-TK cDNA into pLenti-puro (Addgene # 312043). Lentiviral particles were generated by transfection of HEK293FT cells with either LV-mCherry, Lenti-GFP or LV-TK plus pLP1, pLP2 and pLP-VSVG (the latter three from Invitrogen, Walham, MA, USA) using the Mirus transfection reagent (Thermo Fisher). Viral particles were collected 24 and 48 h after transfection, concentrated using vivaspin centrifugation columns (3.000 MWCO, Sartorius, Göttingen, Germany) and were stored at −80°C for further use.

### 2.5. Identification of matrix-metalloproteinase expression (MMP) and activity

MMP expression was analyzed by immunoblot in conditioned medium of LX-2 cells, generated by cultivating the cells in serum deprived medium for 48 h, followed by protein concentration using vivaspin centrifugal concentrators columns (MWCO 3.0 kD, Sartorius GmbH, Göttingen, Germany). Protein contents were analyzed according to Bradford using Rotiquant (Roth, Karlsruhe, Germany). Either 20 or 40 μg of protein was loaded on 10 % polyacrylamid gels and subsequently blotted on PVDF membranes. For the detection of specific proteins, the following antibodies were used: anti-MMP-2 (AF902, 2 μg/ml; R&D Systems, Abington, UK); anti-MMP-9 (MAB-911, 4 μg/ml, R&D); MMP-7 (IMG-3873; 0.5 μg/ml, IMGENEX); MT1-MMP/MMP-14 (D1E4, 1:1000, Epitomics); TIMP-2 (MAB971, 1:500, R&D). Protein contents were visualized on a ChemiDoc MP system using the ImageLab software for quantification (both from Bio-Rad Laboratories GmbH, Munich, Germany). For the demonstration of equal loading, the membranes were subsequently stained with Ponceau S (Sigma-Aldrich). MMP-2 and −9 activity was determined using gelatine containing zymography gels (Thermo Fisher Scientific, Karlsbad, CA, USA) as previously described (37).

### 2.6. RNA Sequencing and bioinformatics data analysis

For RNA sequencing, 2 × 10^6^ cells were seeded, allowed to grow over night, washed twice with ice cold PBS and collected by trypsination. Cell pellets were snap-frozen, stored at −80°C and sent for RNASeq to IMGM Laboratories (Martinsried, Germany). The RNA library was prepared using the NEBNext® Ultra II Directional mRNA Library Prep Kit (New England Biolabs, Frankfurt/Main, Germany). The quality of RNA and the library was determined by Nanodrop and Qubit (both Thermofisher). Sequencing was performed on a NovaSeq6000 next generation sequencing 75 SR system (Illumina, San Diego, CA, USA). Bioinformatic RNASeq analysis was done using the CLC Genomics Workbench v.21.0.5 (Qiagen, Hilden Germany). Signal processing and de-multiplexing was performed using bcl2fastq 2.20.0.422 tool. CLC Genomics Workbench v.21.0.5 (Qiagen, Hilden Germany) was used to length, quality and adapter trimming and mapping reads to GRCh38 reference genome. Gene level read counts were calculated using htseq-count tool (HTSeq 0.11.1) based on GeneCode annotation (version 42, freeze 04.2022). Principle component analysis (PCA) plots and correlation heatmaps were generated with raw and library normalized counts. Visualization of transcriptome landscape was performed with oposSOM R package that allows for dimension reduction and intuitive visualization of over- and underexpression of co-expressed gene clusters (38). Differential gene expression analysis was performed using Deseq2 (39) and edgeR (40) R packages. Since it is known that different pipelines for differential expression analysis can produce different results (41) and to increase robustness of the analysis we confined our analysis to genes that were considered as differentially expressed (logFC > |0.5|, p_adj_ = 0.05) by both tools. Mining for enrichment of migration-related gene sets from Gene Ontology (42) was performed using one-tailed Fishers’ exact test. Gene sets with FDR adjusted p values < 0.05 were considered significantly enriched in differentially expressed gene lists.

### 2.7. Adenoviral vectors and infection

The oncolytic adenovirus subtype 5 XVir-N-31 (also named Ad-Delo3-RGD) has been described in detail in (27-29). Infectious virus particles were generated in HEK293^Mx^ cells, purified and titrated as described in detail in (29). For the determination of virus production, 300.000 shuttle cells were infected with increasing moieties of infection (MOI) of XVir-N-31. Oncolysis was firstly determined microscopically by observing virus-replication associated cytopathic effects (CPE). At optimal oncolysis (48 - 72 h after infection) the supernatants of infected cells were collected and infectious virus particles were measured using the Adeno X Rapid Titration Kit (Takara Bio Europe, Saint-Germain-en-Laye, France).

### 2.8. Immunostainings and immunofluorescence

Hexon staining was performed using the Adeno X Rapid Titer Kit (Takara Bio Europe, Saint-Germain-en-Laye, France). Immunofluorescence was performed using the YB-1 antibody from Santa Cruz (#R0409, Santa Cruz Biotechnology, Heidelberg, Germany). As a secondary antibody anti-Mouse IgG Alexa Fluor™ Plus 680 (Invitrogen) was used. Nuclei were counterstained using 4’,6-Diamidino-2-phenylindol containing mounting medium (Vectashield, Biozol Diagnostica GmbH, Eching, Germany). Fluorescence was analysed using a Zeiss LSM 710 confocal microscope (Carl Zeiss AG, Oberkochen, Germany).

### 2.9. Animal experiments

Animal work was performed in accordance with the German Animal Welfare Act and its guidelines (e.g. 3R principle) and was approved by the regional council of Tübingen (approval N02/20G). NOD.Cg-*Prkdc*^*scid*^ *Il2rg*^*tm1Wjl*^/SzJ (NSG) mice (Jackson Laboratory, Maine, USA) were bred in IVC cages in the animal facility of the institute under sterile conditions and used at an age of 2 - 6 months. INA was performed as described (43) using 24 μl PBS containing up to 4 × 10^6^ cells. For pathology, the mice (n=3 per group) intranasally received either PBS or 4 × 10^6^ LX-2 FR cells and were sacrificed 3 months later. Organs were collected, fixed in formalin and paraffin-embedded (lungs and trachea), or were fixed in 4% PFA (brains), stained with hematoxylin/eosin (HE; Sigma-Aldrich) and subjected to pathological analyses by an experienced pathologist from the Department of Mouse Pathology (Institute for Pathology, University of Tübingen, Germany). All samples were scanned with the Ventana DP200 (Roche, Basel, Switzerland) and processed with the Image Viewer MFC Application. Final image preparation was performed with Adobe Photoshop CS6. For the detection of olfactory system and brain infiltrating shuttle cells, 4 × 10^6^ PAR, FR or FR/TK cells were intranasally applied. Mice were sacrificed 6, 15 and 30 h later. Brains and olfactory tracts of mice were prepared in parallel, frozen on dry ice and sectioned (10 μm). For the determination of tumor tropism, the mice received an intrastriatal, stereotactical guided injection of 100.000 LN-229^GFP^ cells. Twentyone days later, INA of shuttle cells was performed as described before. Images as well as brain tile scans were processed using a Leica DMi8 microscope (Leica, Wetzlar, Germany). Shuttle cells in the olfactory epithelium, olfactory bulb, hippocampus, striatum, thalamus and cerebral cortex were quantified using Image J.

### 2.10. Statistics

All *in vitro* experiments were performed at least thrice if not mentioned otherwise. For *in vivo* experiments, the group and sample size are indicated in the figure legends. Statistical analyses were done with two-tailed Student’s t-test, ANOVA or Wilcoxon Test using GraphPad Prism 9.5 (GraphPad Inc., San Diego, CA, USA). The results are represented as mean ± standard error mean (SEM). p-values of < 0.05 are considered as statistically significant (ns: not significant; * p < 0.05; ** p < 0.01; *** p < 0.001; **** p < 0.0001).

## 3. Results

### 3.1. Generation of optimized LX-2 shuttle cells

To develop an optimized shuttle cell line that can be tracked in vivo after INA, we generated LX-2 cells expressing mCherry (PAR). By serial selection cycles for fast migration, we subsequently isolated an offspring cell line with enhanced cell motility that we called „fast runner” (FR). In addition, FR cells were armed, by lentiviral transduction, to express the suicide prodrug gene HSV-TK as a safety switch (FR/TK) which can be used for their elimination by Ganciclovir (GCV). Both FR and FR/TK cells showed an elevated migratory capacity *in vitro* (Fig. 1A). Using live cell imaging we determined the velocity of cells during their movement in culture over a period of 24h. Both, the speed as well as the overall (accumulated) distance the cells migrated during the fixed period time was significantly higher in FR cells compared to PAR cells (Suppl. Fig. 1A). In addition, the FR and FR/TK cell’s capability to migrate through a matrigel layer was at least double as that of PAR cells (Fig. 1B, Suppl. Fig. 1B). The elevated cell motility was not an effect of elevated cell proliferation of FR or FR/TK cells as the doubling time of these cells (∼ 98h) only marginally differ from that of PAR cells (∼118 h; Suppl. Fig. 2). This difference should not have any effect in the short time migration assays we performed. Finally, genomic integration of the HSV-TK gene into FR cells rendered them highly vulnerable to GCV (EC_50_ FR: ∼ 767 μM, EC_50_ FR/TK: ∼ 45 μM; Suppl. Fig. 3).

**Figure 1:**
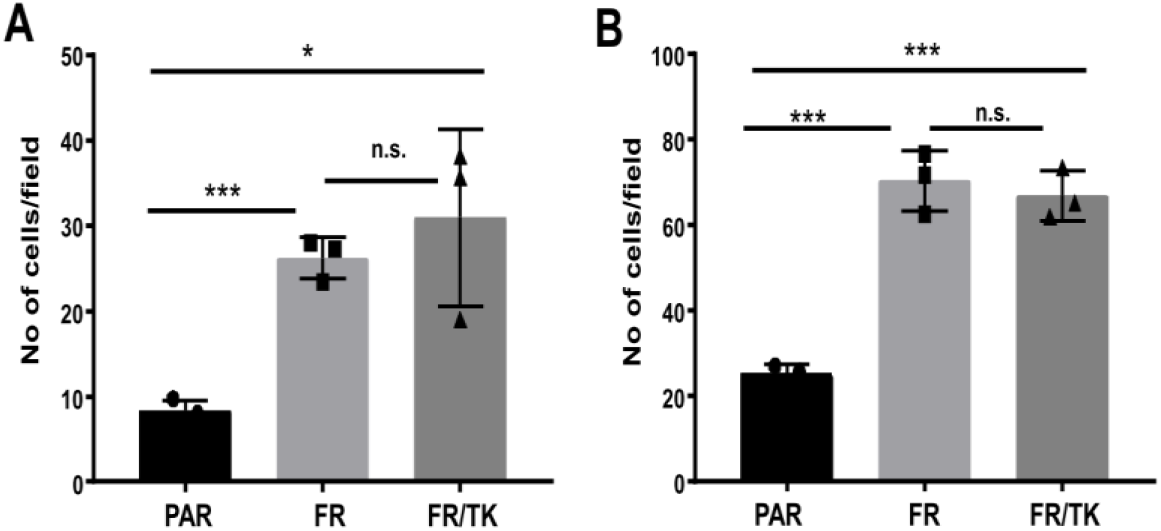
Motility of PAR, FR and FR/TK cells in vitro. A. Migration assay of PAR, FR and FR/TK cells. Cell migration was performed for 5 h using conditioned medium from LN-229 GBM cells as an attraction medium B. Invasion assay. The cells were seeded as in A and invasion was performed for 6 h (A/B: n=3, SEM, t-test, n.s.: not significant; * p <0.05, *** p<0.001).

We further investigated the migration capability of the FR and FR/TK cells *in vivo* after INA. FR and FR/TK cells had migrated significantly faster than their PAR counterparts to the olfactory bulb (Fig. 2A/B) 6 h post INA. Elevated numbers of FR and FR/TK cells were also observed in the cerebral cortex, striatum, thalamus and hippocampus 6 h post INA. With time, the amount of shuttle cells in the different brain compartments further increased, with a clear significant migration advantage for FR and FR/TK cells (Fig 2C/D, Suppl. Fig. 4).

**Figure 2:**
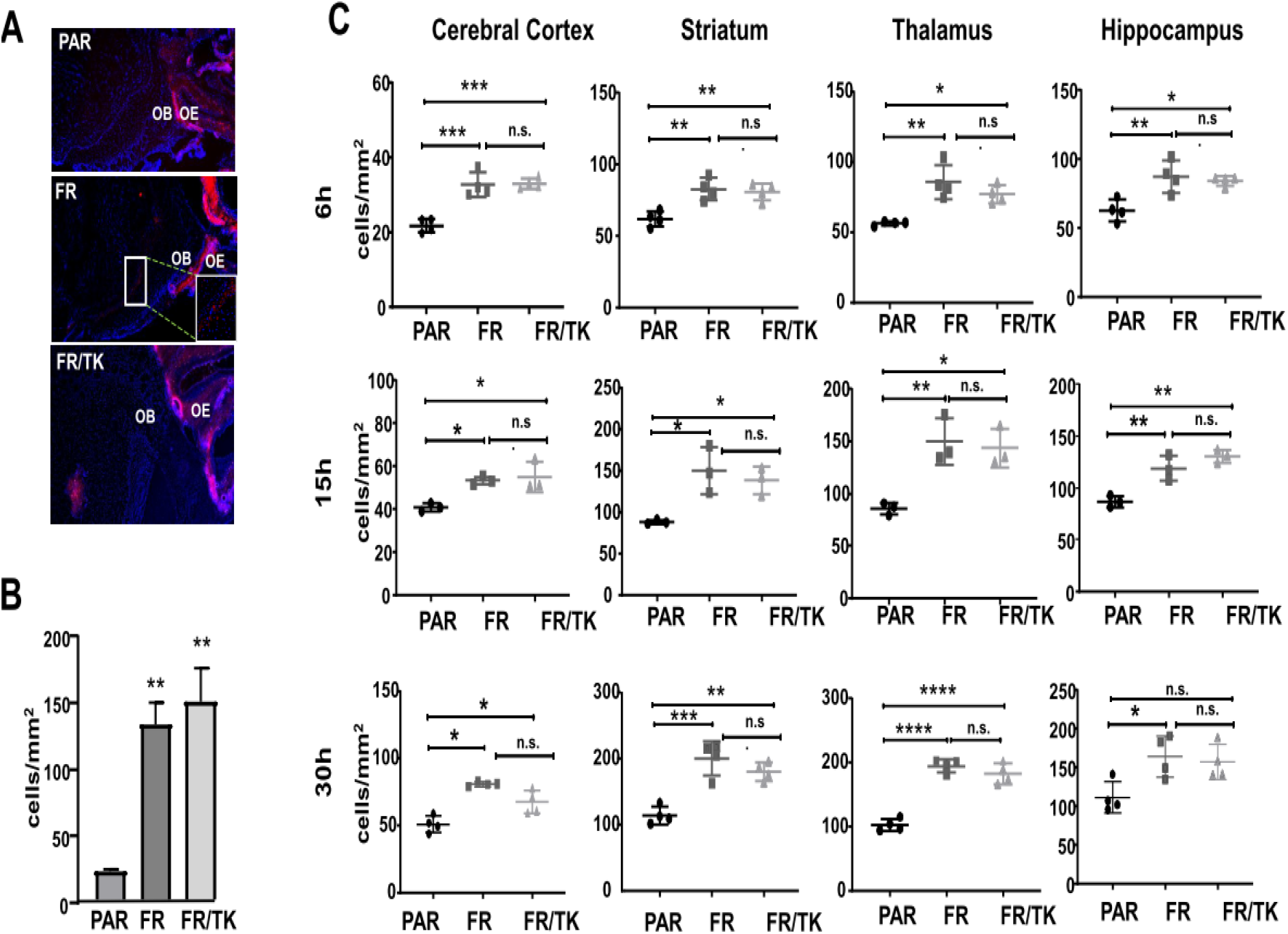
Motility of PAR, FTR and FR/TK cells in vivo. NSG mice intranasally received 4 million shuttle cells. Mice were sacrificed after 6, 15 and 30 h and the amount of mCherry-positive shuttle cells was quantified A. Microphotographs of migrated shuttle cells in the olfactory bulb of the mice 6 h after INA, one picture is exemplarily shown (OE: Olfactory endothelium; OB: olfactory bulb). B. Quantification of migrated shuttle cells in the olfactory bulb as shown in A (3 to 4 mice per group and 3 to 6 slices per mouse were used for quantification; SEM, ANOVA, ** p<0.001) C. Detection of shuttle cells in different mouse brain areas 6, 15 and 30 h after INA (n=3-4 mice per group, 6-8 slices per mouse and brain area were quantified).

We also analyzed whether LX2 cells provide tropism towards GBM cells. For this we firstly performed *in vitro* migration assays using GBM cells or glioma stem cells (GCSs) as attractants seeded in the bottom part of a migration chamber. However, we observed no differences in the cell’s migration capacity towards tumor cells compared to conditioned medium (Fig. 3A). We then traced FR cells post INA in our LN-229^GFP^ GBM bearing mouse model. FR cells were detected in and around the tumor core including the infiltration zones; nevertheless, there were no clear signs of tropism towards this particular tumor (Fig. 3B).

**Figure 3:**
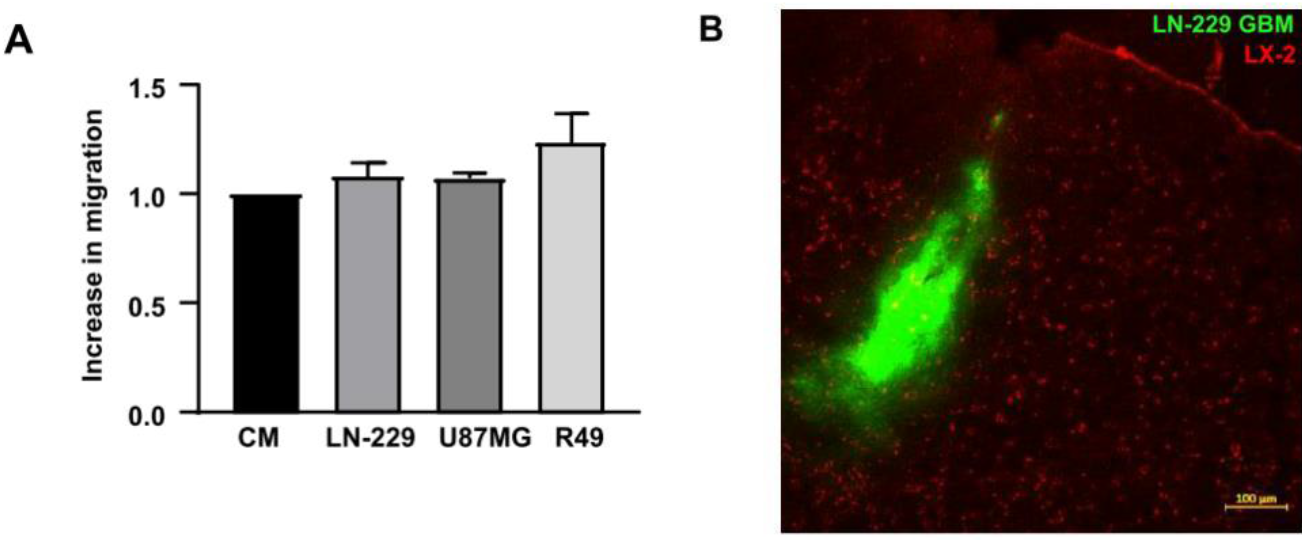
LX-2 PAR cells show no tumor tropism. A. In vitro the migration of PAR cells was performed for 4 h using conditioned medium of PAR cells as attraction medium and serves as a control, or against 200.000 LN-229 or U87MG GBM, or R49 GCSs seeded in the bottom well of a migration chamber as described in the material and methods part (n=2-3, SD). B. LN-229^GFP^ bearing NSG mice received INA of 4×10^6^ FR cells. Microphotographs were taken 72 h later (n=3, one representative picture is shown).

### 3.2. The intranasal delivery of LX-2 cells does not induce pathological changes long time after administration

As LX-2 cells are immortalized cells with a variety of karyotypic alterations (44), we investigated whether their intranasal delivery induces pathological changes and/or might lead to tumorigenesis s in the organs the cells can reach by INA such as the respiratory tract and/or the brain. For this, we intranasally applied either PBS or 4 × 10^6^ FR cells into NSG mice and performed histopathological analysis on the lungs, trachea and brains of these mice 3 months later. All examined organs, both of control and experimental animals, showed normal histology. No acute inflammation was detected in any sample. Five out of 6 animals showed a mild focal presence of macrophages with abundant small lipid vacuoles in the cytoplasm, independent from the treatment group. Finally, none of the animals developed tumors in lungs, tracheas or brains (Fig. 4).

**Figure 4:**
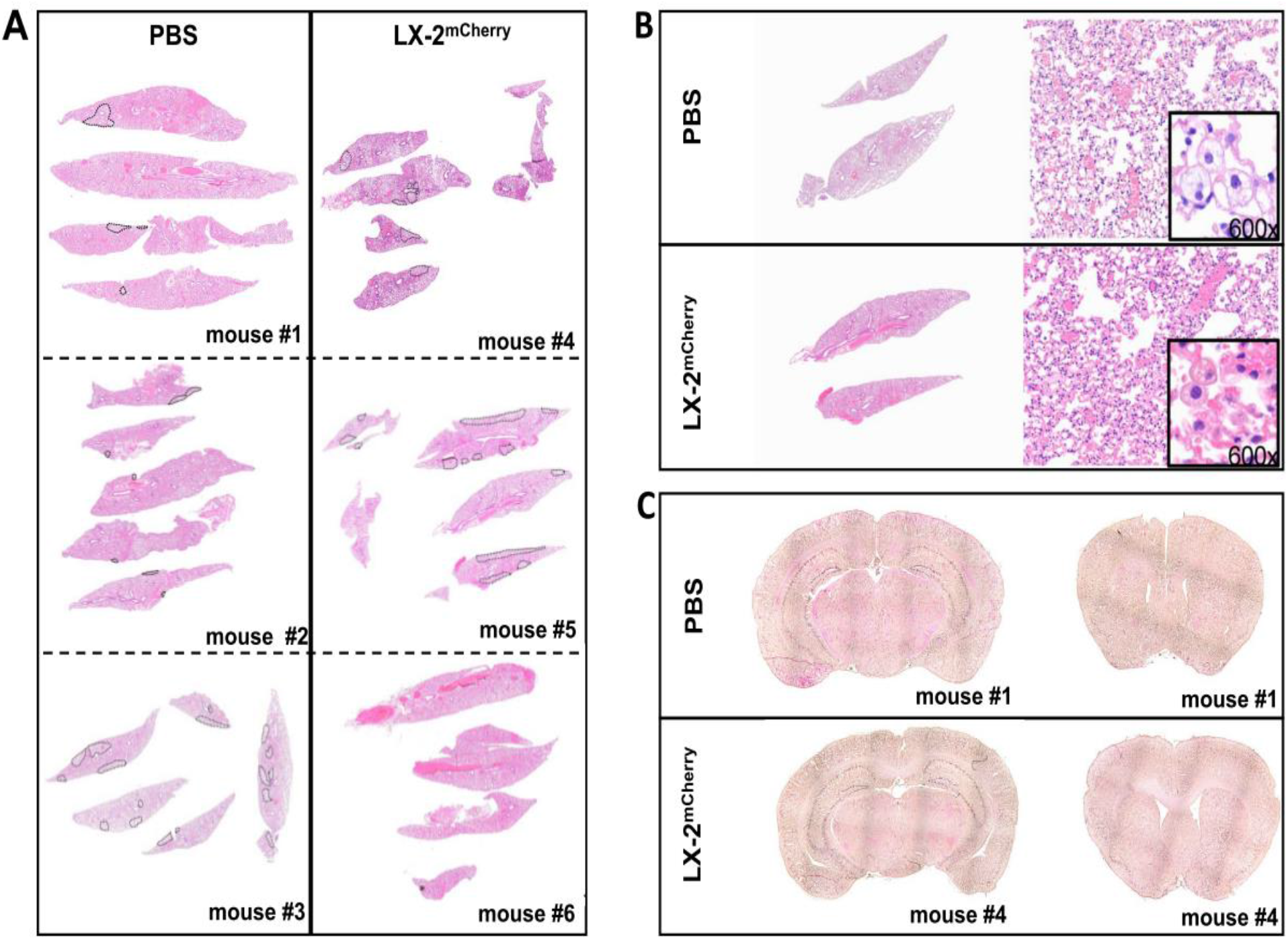
Pathology of mice receiving INA of LX-2 cells. The mice (n=3 per group) intranasally received either PBS or 4 × 10^6^ PAR cells. After 3 months the mice were sacrificed and the tissues were examined. A/B. Histology of lungs and tracheas. All lungs showed abundant foamy macrophages. C. Exemplarily the histology of two different brain areas (coronal sections from the front and middle part of the brain) of one control mouse and one LX-2 receiving mouse is shown.

### 3.3. Molecular characterization of optimized LX-2 shuttle cells

As LX-2 cells are activated HepSCs, they express a variety of matrix remodelling genes like matrix metalloproteinases (MMP)-2, MMP-9, MMP-14/MT1-MMP, as well as tissue inhibitors of MMPs like TIMP-1 and −2 (19, 45). We therefore determined whether these genes were differentially expressed in FR and FR/TK cells compared to PAR cells. Indeed, elevated cell motility in FR and FR/TK cells was accompanied by a slightly elevated MMP-2-, an up to 60-fold elevated MMP-9, and an up to 3-fold MMP14/MT1-MMP protein expression, whilst the tissue inhibitor of metalloproteinase (TIMP)-2 was downregulated. The expression of MMP-7 was not altered (Fig. 5A/B) whereas MMP-1 or MMP-10 proteins could not be detected in any of the cell lines (data not shown). Gelatine-Zymography also showed an elevated activity of MMP-2 and −9 (Fig. 5C). If being significant, mRNA expression of these genes was mainly in concordance with protein expression (Fig. 5D).

**Figure. 5:**
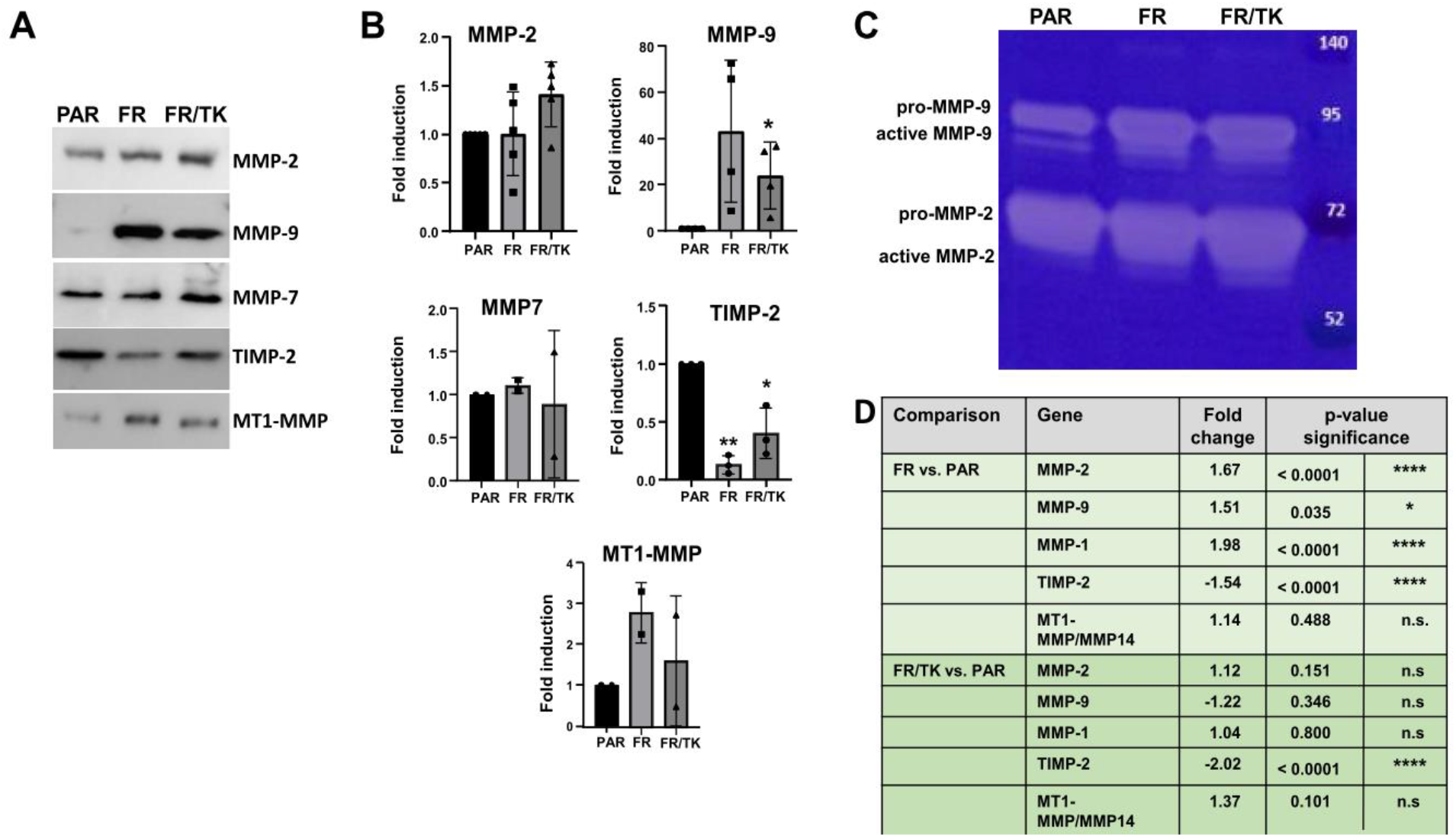
Expression of matrix remodelling proteins. A. Immunoblots showing MMPs and TIMP-2 expression (n=2-4, representative blots are shown). B. Quantification of protein expression (n=2-4, SEM, * p<0.05, ** p<0.01). C. MMP activity in cell supernatants determined by gelatin-zymography (n=3, one representative zymogram is shown). D. RNA expression of MMPs and TIMP-2 in PAR, FR and FR/TK cells determined by RNASeq analysis (n=3, ANOVA; ** p<0.01, **** p< 0.0001).

By selection for FR cells we generated a cell line with an expression transcriptome profile that was clearly separated from those of PAR cells, but was only slightly altered by the introduction of the HSV-TK gene (Fig. 6).

**Figure 6:**
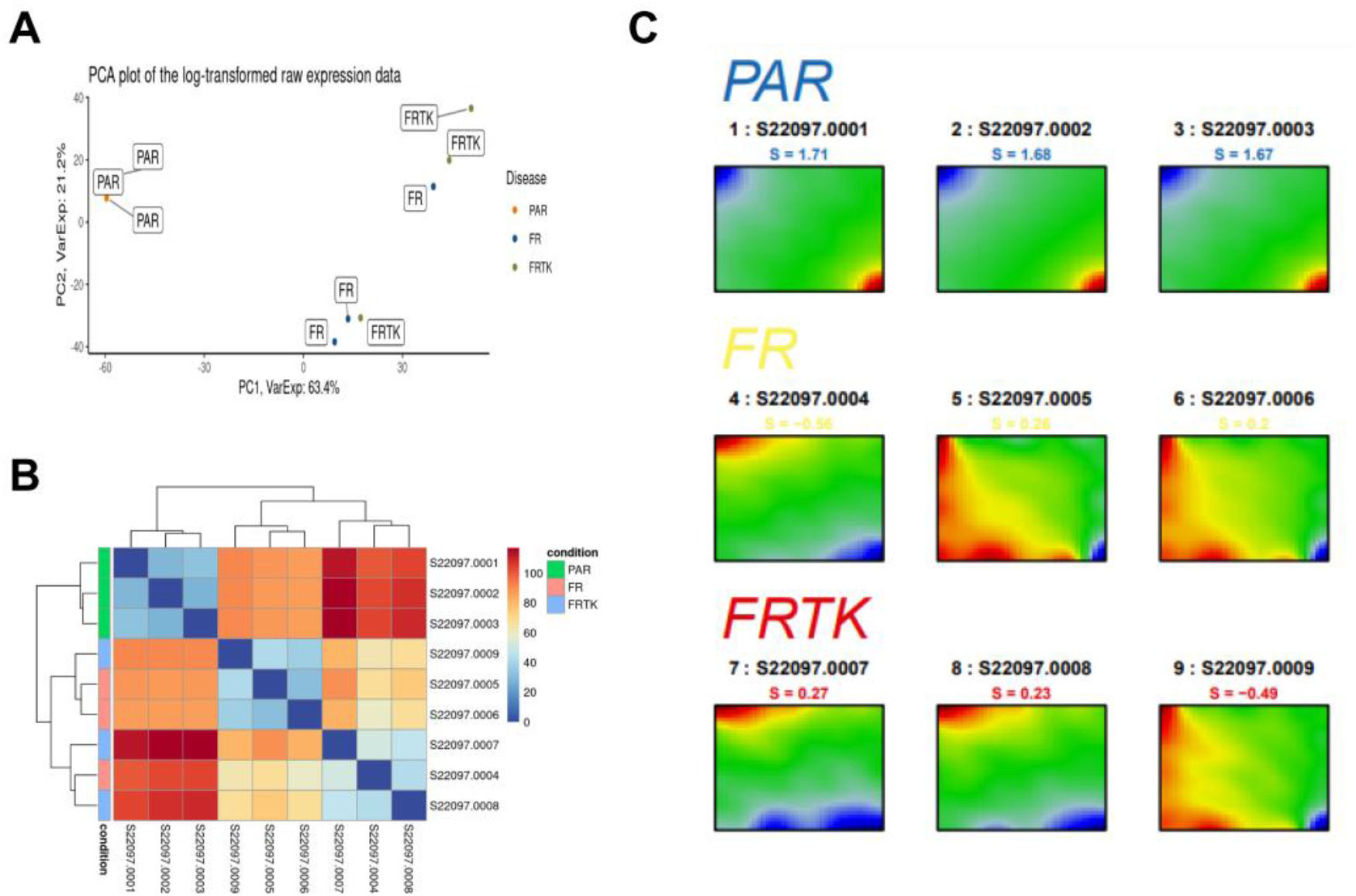
FR and FR/TK represent a group of cells separated from their ancestor LX-2 cells. A. Principal component (PCA) analysis of RNASeq data demonstrates that FR and FR/TK cells present cell lines that can be separated from PAR cells by gene expression (n=3). B. Heat map analysis of PAR, FR and FR/TK cells. C. Self-organizing map (SOM) analysis demonstrates the differences in the transcriptome between PAR, FR, and FR/TK cells (n=3).

A majority of the differentially regulated genes in FR and FR/TK cells compared to PAR cells are involved in the regulation and organization of the cytoskeleton like Mesenchyme Homeobox (MEOX)-2 gene, of genes regulating cell cell-cell adhesions like the P53 Apoptosis Effector Related to PMP22 (PERP) or A Disintegrin And Metalloproteinase (ADAM)-23, of cell polarity like Desmoglein (DSG)-2 and of cell junctions like the Cytohesin 1 Interacting Protein (CYTIP). Additionally, a further altered expression of cell adhesion associated genes popped up in FR compared to FR/TK cells like the teneurin transmembrane protein (TENM). Besides, genes involved in metabolic processes like the Solute Carrier Family 1 Member (SLCA)-3, Glycerolphosphat-O-Acyltransferase (GPAT3) or genes that are known to be expressed in neural cells like the Neural Cell Adhesion Molecule 2 (NCAM2) that is important for the plasticity of the olfactory system, or the Cholinergic Receptor Muscarinic (CHRM)-3 are differentially expressed in FR/TK compared to FR cells (Fig. 7A/B, Table 1).

**Table 1:** Most prominent differentially regulated genes in FR or FR/TK cells vs. PAR cells.

| Gene | Protein | Log<sub>2</sub> fold change PAR vs. FR | Log<sub>2</sub> fold change FR/TK vs. PAR | Function | Reference |
| --- | --- | --- | --- | --- | --- |
| AEBP1 | Adipocyte enhancer-binding protein 1 | ↓<br>-7.08 | ↓<br>-8.41 | Transcriptional repressor. Regulation of proliferation, migration, invasion, EMT and collagen remodelling | Majdalawieh AF et al. (2020) J. Oncol. 2020: 8097872. |
| PERP | p53 apoptosis effector related to PMP22 | ↓<br>-4.91 | ↓<br>-5.5 | Plasma membrane protein. Effects cell-cell adhesion and gene expression. | Jheon AH J (2011) Cell Sci. 2011;124(5):745-54 |
| CHFR | Checkpoint with forkhead and ring finger domains | ↓<br>-4.70 | ↓<br>-5.72 | E3-ubiquitin ligase, of e.g. HDAC1. Located in nucleus and microtubule. Regulates cell cycle entry into mitosis | Privette LM et al. (2008) Transl. Oncol. 1(2):57-64 |
| ADAM23 | A disintegrin and metallo proteinase 23 | ↓<br>-4.97 | ↓<br>-5.02 | Promotes cell adhesion via interaction with αvβ3 integrin | Cal S et al. (2000) Mol. Biol. Cell 11(4):1457-1469 |
| BEX4 | Brain Expressed X-Linked 4 | ↓<br>-4.70 | ↓<br>-6.56 | May play a role in microtubule deacetylation. Participate in the control of cell cycle progression | Gao W et al. (2016) J. Exp. Clin. Cancer Res. 35: 92 |
| SERINC2 | Serine incorporator 2 | ↓<br>-4.04 | ↓<br>-5.29 | Transmembrane protein. Regulate synthesis of phosphatidylserine and sphingolipids. | Inuzuka M et al. (2005). J Cell Biol 80, (42): 35776-35783 |
| MS4A4A/CD20L1 | Membrane spanning 4-domains A4A | ↑<br>9.92 | ↑<br>6.85 | can regulate cell activation by working as ion channels or by modulating the signaling of other immunoreceptors. Might be involved in lymphocyte migration. | Mattiola I (et al. (2021) Trends Immunol 42(9):764-781. Sanyal R et al. Immunol. Cell. Biol. (2017) 95 (7): 571-648 |
| CYTIP | Cytohesin 1 interacting protein | ↑<br>12.36 | ↑<br>12.40 | Sequesters Cytohesin-1 in the cytoplasm and thereby limits its interaction with β2 integrins, thereby reducing cell adhesion. | Heufler C et al. (2008) Immunobiol. 213:729–732 |
| MEOX2 | Mesenchyme Homeobox 2 | ↑<br>7.32 | ↑<br>7.41 | In glioma induces cell proliferation, motility, EMT, focal adhesion formation, and F-actin assembly | Wang J et al. (2022) Cell Death Disease 13: 360 |
| SFRP2 | Secreted Frizzled Related Protein 2 | ↑<br>12.17 | ↑<br>12.53 | Acts as a soluble modulator of Wnt signaling. In osteosarcoma promotes cell invasion | Techavichit P al. (2016) BMC cancer, 16(1): 869 |
| MS4A6E | Membrane Spanning 4-Domains A6E | ↑<br>10.40 | ↑<br>9.79 | Tetraspanin-like protein. Associated with the risk and onset of Alzheimers disease. | Karch, C et al. (2015) Biological psychiatry, 77(1), 43–51. |
| PLA2G4A | Phospholipase A2 Group IVA | ↓<br>-3.98 | --- | Catalyzes hydrolysis of membrane phospholipids to release arachidonic acid which is subsequently metabolized into eicosanoids. Involved in lipid metabolism. | Adler DH et al. (2008) J. Clin. Invest. 118: 2121-2131 |
| MYO1D | Myosin 1D | ↓<br>-5.29 | ↓<br>-5.54 | Unconventional myosin holding EGFR family members at the plasma membrane. | Yoo-Seung K et al. (2019) *Oncogene* 38:7416–7432 |
| ABCA3 | ATP binding cassette subfamily A member 3 | ↓<br>-5.40 | ↓<br>-11.85 | Membrane-associated protein. Member of the superfamily of ATP-binding cassette (ABC) transporters, also functions as a lipid transporter. | Ban N et al. (2007) *Membrane Transport, Structure & Function* 282(13): 9628-9634 |
| NXN | Nucleoredoxin | ↓<br>-5.31 | ↓<br>-4.45 | Suppresses metastasis of hepatocellular carcinoma by downregulation of the EMT protein SNAIL. | Zhang Y et al. (2022) *Cell Death Dis* 13(8): 676 |
| MAGEA3 | MAGE Family Member A3 | ↑<br>7.76 | ↑<br>7.41 | Depletion of MAGEA3 resulted in reduced cell proliferation and increased apoptosis upon growth factor deprivation. | Biswajit D et al. (2019) *J Exp Clin Cancer Res* 38, 294 |
| PDGFRA | platelet-derived growth factor receptor alpha | ↓<br>-3.95 | ↓<br>-5.11 | Receptor tyrosine kinase. Signaling pathways stimulated by PDGFRA control many cellular processes such as cell growth, proliferation and survival. | Cao Y (2013) *Trends Mol Med* (2013). 19, 8:460-473 |
| DSG2 | Desmoglein 2 | ↓<br>-3.15 | ↓<br>-4.79 | Localized to desmosome structures at regions of cell-cell contact. Structurally adheres adjacent cells. | Ebert LM (2016) *Angiogenesis* 19: 463–486 |
| PRXL2A | Peroxiredoxin Like 2A | --- | ↓<br>-4.41 | Involved in redox regulation of the cell. | Guo F et al. (2015) *Biochem J*;472(3): 309-318 |
| NCAM2 | Neural Cell Adhesion Molecule 2 | --- | ↓<br>-4.60 | Cell adhesion molecule. Controls neuronal migration. | Parcerisas A (2021) *Int J Mol Sci.* 22(18): 10021 |
| GATA3 | GATA3 | ↑<br>9.88 | ↑<br>11.42 | Zinc finger transcription factor. Involved in HepSC activation. | Lin A et al. (2022) *Phytomedicine* 103:154199 |

**Figure 7:**
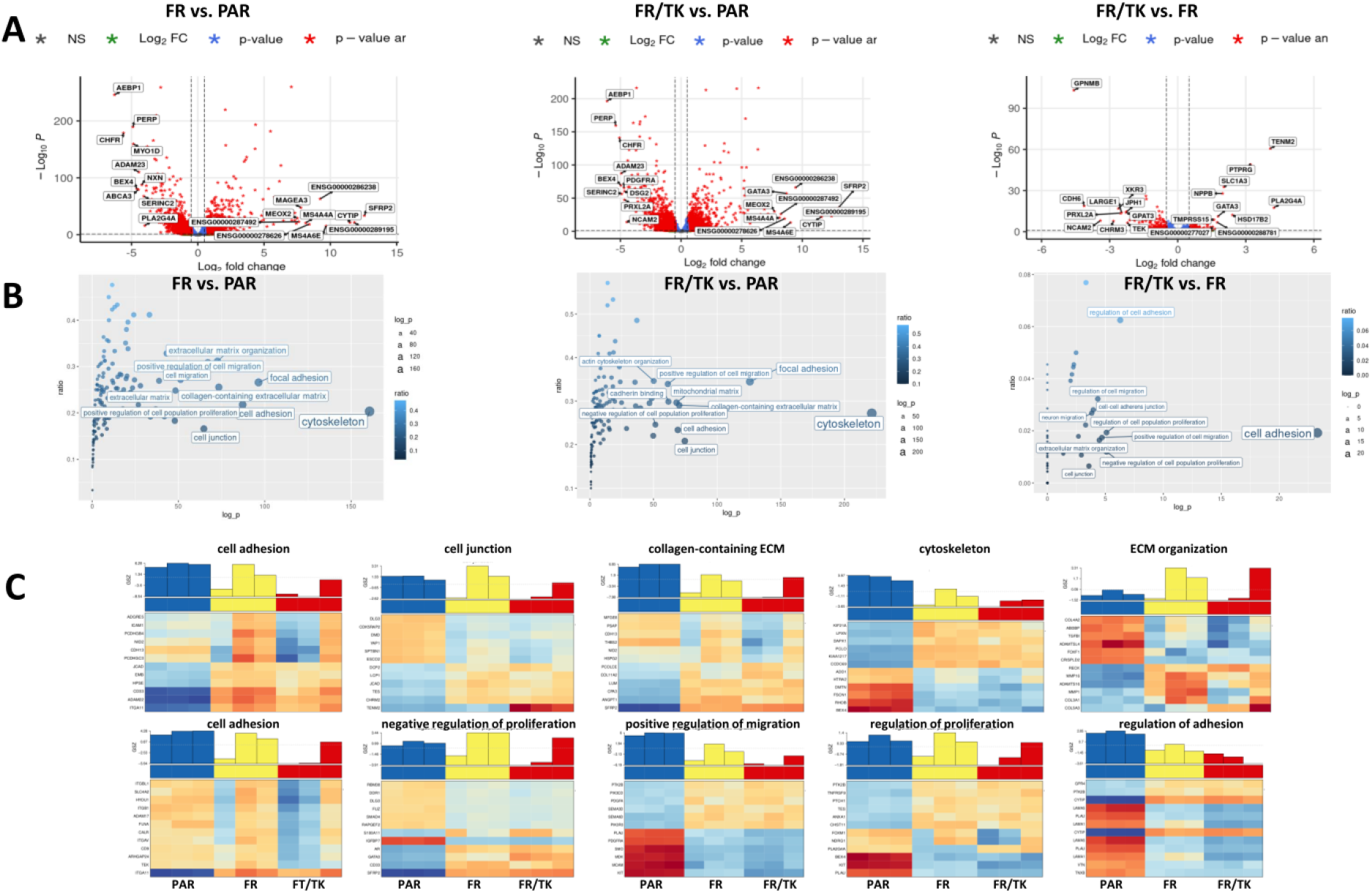
FR and FR/TK represent a group of cells separated from their ancestor LX-2 cells. A. Volcano plots of the most prominent differentially regulated genes in PAR, FR, and FR/TK cells. B. Pattern hunter analysis demonstrates the differences in cellular processes in PAR, FR, and FRTK cells. C. Heatmap of gene expression and their association to cellular processes.

Further heatmap analyses indicate that especially genes that are part of the cytoskeleton, that positively regulate cell migration or that are either involved in the process of cell adhesion or that are regulators of cell adhesion were downregulated in FR and FR/TK cells. Additionally, the expression of several genes negatively regulating cell proliferation were differentially expressed in FR and FR/TK cells which complies with the elevated proliferation rate of FR and FR/TK cells we observed (Fig. 7C, Suppl. Fig. 2).

### 3.4. XVir-N-31 efficiently replicates in LX-2 cells

Our goal was to generate an optimal shuttle cell line that can be used to efficiently transport therapeutic agents form the nose into the brain. In a proof of principle concept, we used as a cargo the OV XVir-N-31 that has been shown to efficiently eliminate GBM if injected intratumorally. As the replication of XVir-N31 is dependent on YB-1, a protein that is highly expressed in most immortalized cell lines and in high grade GBM, but not in non-neoplastic brain cells (46), we firstly determined YB-1 expression in LX-2 cells and could demonstrate that YB-1 is expressed in these cells (Fig. 8).

**Figure 8:**
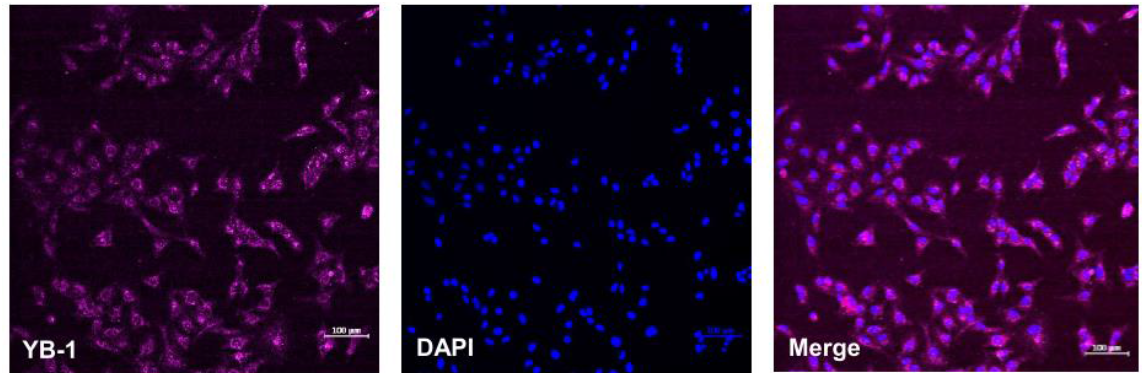
LX-2 cells express YB-1. Immunofluorescence staining of YB-1 (left). In addition, cells were stained with DAPI to visualize nuclei (middle and right; bars = 100 μm).

Thereupon, we analysed XVir-N31 replication in LX-2 cells and in mouse HepSCs and compared it to replication in HB1.F3 human NSCs, a cell line that has been previously used as shuttle cells for OAVs (15). A virus replication dependent cytopathic effect (CPE) in LX-2 cells was observed earliest 48 h after infection, with an optimum around 72h (Fig. 9A). In contrast to LX-2 cells, CPE and hence a virus mediated cell killing is much faster in NSCs, rendering most NSCs dead after 48h and all around 60 h after infection. The observed CPE was accompanied by the production of the adenoviral hexon capsid protein in infected cells (Fig. 9B) and the production and release of infectious virus particles into the medium. However, even if the production of infectious XVir-N-31 particles did not significantly differ between LX-2 and HB1.F3 cells when supernatants were collected at 48h after infection (Fig. 9C), the delayed virus-mediated killing of LX-2 cells might give them more time to migrate through the brain towards the tumor if OV-loaded LX-2 cells are applied intranasally. We also tested mouse M1-4HSV HepSC for virus production. However, even if M1-4HSCs can be easily infected with and killed by XVir-N-31 as shown by CPE and hexon staining, production of XVir-N-31 infectious particles by these cells as with many other mouse cells are negligible (Fig. 9C).

**Figure 9:**
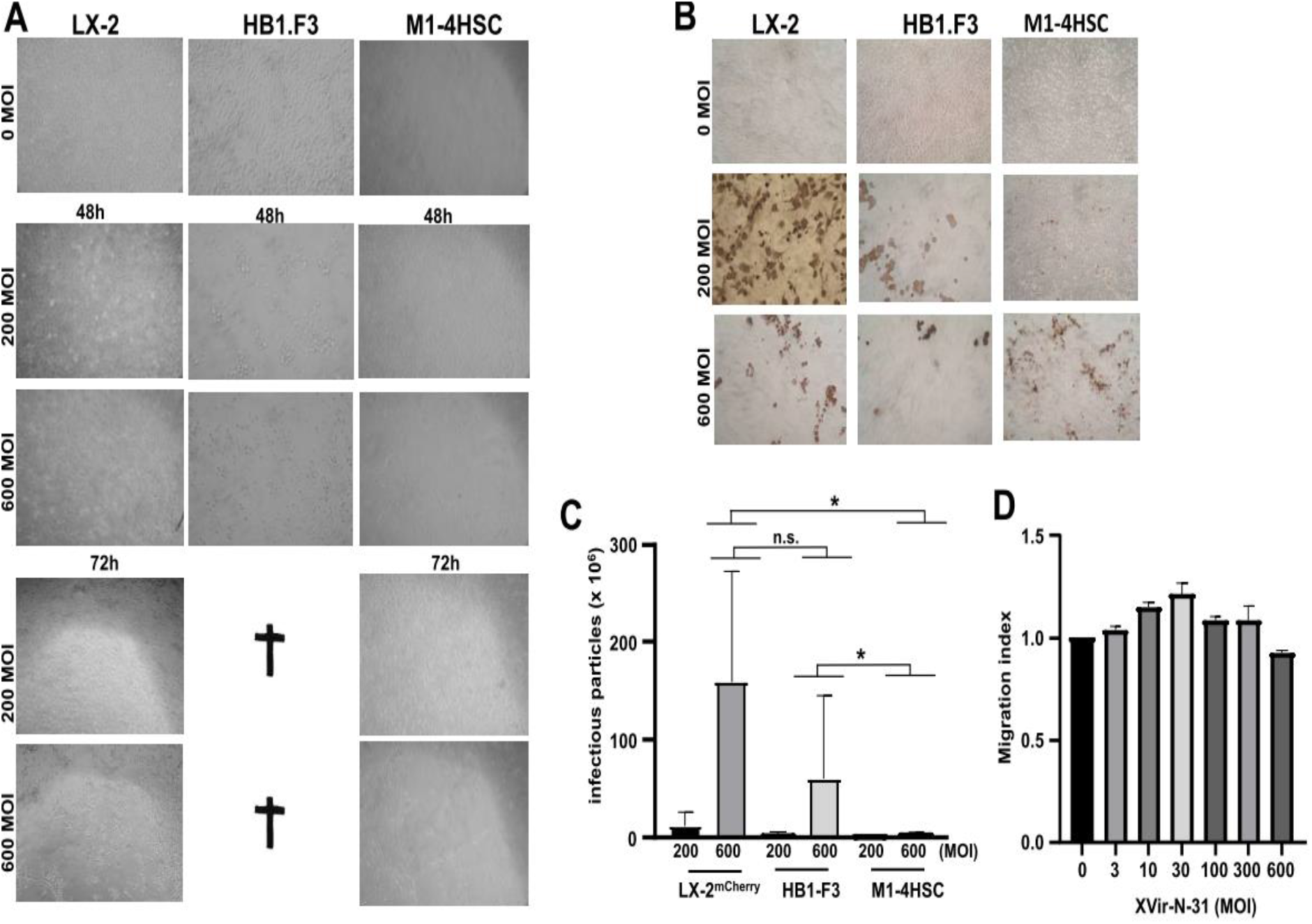
LX-2 cells can be used to produce the oncolytic adenovirus XVir-N-31. A. Photographs of XVir-N-31 infected LX-2, HB1.F3, and M1-4HSCs. The cells were infected with either 200 or 600 MOI or were left untreated. 48 and 72 h after infection pictures were taken to show the XVir-N-31 mediated cytopathic effect (CPE; n= 3-7). Notice that HB1.F3 cells were completely detached and dead 72 h post infection. B. The cells were infected as in A. 48 h (HB1.F3) or 72 h (LX-2, M1-4HSC) later the cells were stained for the adenoviral hexon capsid protein (n=3, one representative picture is shown). C. XVir-N-31 infectious particle production. 300.000 cells were infected with the indicated MOI of XVir-N-31. 48h later supernatants from infected cells were collected and infectious virus particles were determined (n=2-6, SEM, * p<0.05; ANOVA). D. Migration of XVir-N-31-infected LX-2 cells. 100.000 cells were infected with the indicated MOI. After 2 h, the cells were intensively washed with PBS and a Boyden chamber migration assay was performed. After 5 h migrated cells on the bottom side of the membrane were collected and counted using CTB (n=2-7, SEM, ANOVA/ Wilcoxon test).

## 4. Discussion

Our data identifies LX-2 cells as a robust cell line that can be used as a shuttle to transport therapeutic agents into the brain. In contrast to MSCs that have been demonstrated to provide tumorigenicity (7, 8) or to HB1.F3 NSCs that have been reported, at least in immune-compromized mice, to form a tumor mass in the lungs after INA (6). Therefore, a stem cell-based approach generally had certain drawbacks regarding safety. Thus, we needed to ensure the safety of our approach. We did not observe any pathological changes or tumor development in the respiratory tract nor in the brains of NSG mice even months after INA of a high amount of LX-2-cells (Fig. 4). Nevertheless, as a safety backup to eliminate intranasally applied cells, we introduced the HSV-Thymidin kinase gene into LX-2 leading to cell suicide by oral administration of GCV, in case of adverse events. HSV-TK incorporation in these cells raises the vulnerability towards GCV at least 15-fold (Suppl. Fig. 1). This safety switch might be obsolete when using LX-2 cells as vehicles for OVs such as XVir-N-31, since virus replication ensures the lysis of all virus-loaded shuttle cells (Fig. 9). However, the HSV-TK expression in LX-2 cells opens the possibility of using LX-2 cells as vehicles for neuroprotective agents to treat neurodegenerative diseases.

Unexpectedly, we observed no tropism of INA applied LX-2 cells, neither to specific brain regions nor to GBM (Figs. 2, 3). This seemingly disadvantageous feature of LX-2 may turn to be beneficial regarding the treatment of neurological diseases where multiple disease loci are distributed throughout the brain, as it occurs for instance in neurodegenerative disorders or in infiltratively growing tumors such as GBM. In case of GBM this lack of tropism of LX-2 cells may help target invaded tumor cells located distantly from the tumor core. A tumor tropism of NSCs to GBM has been suggested to be induced by a gradient of the vascular endothelial growth factor (VEGF) as a guidance signal evidenced by in vivo studies in nude mice with orthotopic human GBM xenografts (47). However, it is reasonable to assume that the environment of invaded GBM cells located distantly from the original tumor core lacks hypoxia as a major inducer of VEGF secretion (for review see (48)) leading to angiogenesis. Thus, no VEGF gradient will be present to guide OV-loaded NSCs towards invaded GBM cells. In contrast to NSCs, LX-2 cells do not show GBM tropism and after INA are rather distributed throughout the brain (Fig. 2,3), which suggests a higher likelihood of targeting invaded GBM cells by OV-loaded LX-2 shuttle cells than using intranasal NSCs for this approach.

We further optimized LX-2 for their application by INA using our method of selecting highly migratory population of cells (17) to generate an offspring cell line called “fast runners” (FR). In their transcriptome profile both FR as well as their HSV-TK armed siblings (FR/TK) are strictly separated from the original PAR cells (Fig. 6) which suggest that by the selection process the cells have changed their phenotype and behaviour. Compared to PAR cells, FR cells demonstrated an elevated capacity to migrate in vitro and in vivo which was affected by the expression of HSV-TK (Fig. 1 and 2). The velocity of FR cells was 2.5 x greater compared to PAR cells (Suppl. Fig. 1A). To explore the mechanisms behind the enhanced motility of FR and FR/TK cells we performed further molecular analyses focussing on motility associated factors and RNASeq analyses to identify differentially expressed genes that might be associated with processes involved in cell migration and invasion. Matrix-metalloproteinases (MMPs) are recognized to be common motility genes necessary for invasion (for review see (49)). We found upregulated MMP-2 and −9 in FR and FR/TK cells both on the protein level as well as at their activity status, whereas the MMP-inhibitor TIMP-2 was downregulated (Fig. 5). In addition, other genes that are either directly involved in cell adhesion or its regulation, or are components of the cytoskeleton or those regulating cell motility are differentially expressed in FR and FR/TK cells compared to PAR cells (Fig. 7). Furthermore, as summarized in Table 1, some of differentially regulated genes are more indirectly involved in the regulation of cell motility like the tetraspanin family member membrane spanning 4-domains A4A (MS4A4A/CD20L) that act as ion channels and regulate immune cell activation and lymphocyte migration (50), the zinc finger transcriptional factor GATA3 that is involved in the activation of HepSCs (51) or the mesenchymal homeobox protein MEOX2 that is associated with the assembly of F-actin and induces the process of epithelial to mesenchymal transition (EMT) in cancer (52). In cancer cells, changes in sphingolipid metabolism affects cell motility by mediating cell adhesion (53). In FR and FR/TK cells the Serin Incorporator 2 (SERINC2) that regulates the synthesis of phosphatidylserine and sphingolipids (54) is downregulated 4 - 5 fold in FR and FR/TK cells, suggesting that also changes in the lipid metabolism may be one of the reasons for the elevated motility of FR and FR/TK cells.

As an example of a therapeutic cargo that should be transported to the brain by INA, we used the YB-1 dependent oncolytic adenovirus XVir-N-31 which has been previously reported to prolong the survival of GBM-bearing mice when applied intratumorally (26, 28, 29, 55). The expression of YB-1 in LX-2 assures efficient replication of XVir-N-31 in these cells. However, in comparison to the HB1.F3 NSC cell line, the replication cycle of XVir-N-31 in LX-2 cells is prolonged. In NCS and using the same amount of infectious virus particles as for LX-2 cells, most NSCs were dead 48 h after infection and no viable NSCs were detected further at 72h post infection (Fig. 9). This period is prolonged in LX-2 cells, suggesting that XVir-N-31-loaded LX-2 cells have more time to travel from the nose into the brain. Moreover, the OV-load did not affect the LX-2 cells’ ability to migrate (Fig. 9D). Altogether our results suggest that LX-2 cells are valuable candidates for delivering OVs and potentially other therapeutic agents to the brain via intranasal administration. Further studies assessing the therapeutic efficacy of intranasally delivered XVir-N-31-loaded optimized LX-2 (FR or FR/TK) cells in comparison to the commonly used direct intratumoral injection of the OV in in vivo xenograft models of GBM are underway.

## 5. Conclusions

This study provides the first proof of principle for using a hepatic stellate cell (HepSC) line as a vehicle for oncolytic adenoviruses delivery to the brain via intranasal administration. Our results identified the HepSC line LX-2 as a robust carrier of the oncolytic adenovirus XVir-N-31 with a significantly extended lifetime of virus-carrying cells in comparison to the neural stem cells. The migratory capacity of LX-2 was optimized by a selection procedure allowing the generation of an offspring cell line with improved motility and capacity to reach the brain faster than its parental cell line after intranasal administration. LX-2 applied intranasally has been proven safe at least for the target organ (brain) and for the lung, as a potential locus of unintentional delivery after intranasal administration.

## Abbreviations

AD: Alzheimers Disease
ANOVA: analysis of variance
ADAM23: A Disintegrin And Metalloproteinase 23
bFGF: basic fibroblast growth factor
BSA: bovine serum albumin
CHRM: Cholinergic receptor muscarinic
CM: conditioned medium
CNS: central nerve system
CPE: cytopathic effect
CTB: Cell titer blue
CYTIP: Cytohesin 1 Interacting Protein
DMEM: Dulbecco’s Modified Eagle’s Medium
DSG-2: Desmoglein-2
EC_50_: Median effective concentration
EGF: epidermal growth factor
FC: fold change
FCS: fetal calf serum
FDR: false discovery rate
FR: “fast runner”
GBM: glioblastoma
GFP: Green fluorescence protein
GPAT3: Glycerolphosphat-O-Acyltransferase
HepSC: Hepatic stellate cell
HSV-TK: Herpes Simplex Virus Thymidin Kinase
INA: Intranasal application
IT: intratumoral
KEGG: Kyoto Encyclopedia of Genes and Genomes
o/n: overnight
LIF: Leukemia inhibitory factor
LV: Lentivirus
MEOX2: Mesenchyme Homeobox 2
MMP: Matrix-Metalloproteinase
MSC: Mesenchymal stem cell
NCAM2: Neural Cell Adhesion Molecule 2
NEAA: Non-essential amino acids
NSC: Neural stem cell
OV: Oncolytic virus
OAV: Oncolytic adenovirus
OVT: oncolytic virotherapy
PAR: parental cells
PERP: P53 Apoptosis Effector Related to PMP22
PD: Parkinson’s Disease
P/S: Penicillin/streptomycin
RT: Room temperature
SD: Standard deviation
SEM: Sstandard error of the mean
SLCA-3: Solute Carrier Family 1 Member 3
TENM: Teneurin transmembrane protein
TK: Thymidin kinase
VEGF: Vascular endothelial growth factor

## Data accessibility

The datasets generated during and/or analyzed during the current study are available on the Mendeley data repository as soon as the manuscript has been accepted (DOI: 10.17632/prcbbhyb3z.1). Raw data of microscopy photographs are available per request through the corresponding author. Final data from RNASeq analyses were deposited at GEO (accession number XXXXXXX) and will be available as soon as the manuscript has been accepted.

## Authors Contribution

UN and LD designed the study. UN supervised the study, has full access to all data in the study and take responsibility for the integrity of the data and the accuracy of the data analysis. UN, AEA and LD wrote the manuscript. MS did critical revision of the manuscript. AEA, UN, LD, MK, LM, AA, SH, IGM and LQM performed experiments and analyzed data. PSH and WM provided necessary material.

## Acknowledgment

This work was supported by German Cancer Foundation (grant number 70113907). The authors wish to thank Dr. Marine Buadze and Luisa Scholz for their excellent assistance in tracking LX-2 in vivo and in vitro. MS is supported by the Robert Bosch Stiftung, Stuttgart, Germany.

## Supporting Information

***Supplementary Figure 1:*** A. Cell motility of PAR and FR cells determined by live cell imaging.

***Supplementary Figure 2:*** Proliferation of PAR, FR and FR/TK cells.

***Supplementary Figure 3:*** GVC vulnerability of FR and FR/TK cells.

***Supplementary Figure 4:*** Detection of shuttle cells in different brain area of NSG mice.

***Supplementary Figure 5:*** Uncropped immunoblots as shown in partial in Figure 5.

***Supplementary Figure 6:*** Cell line authentication data.

