## Supplementary material for "Development of an optimized, non-stem cell line for intranasal delivery of therapeutic cargo to the central nervous system": Suppl. Fig 1

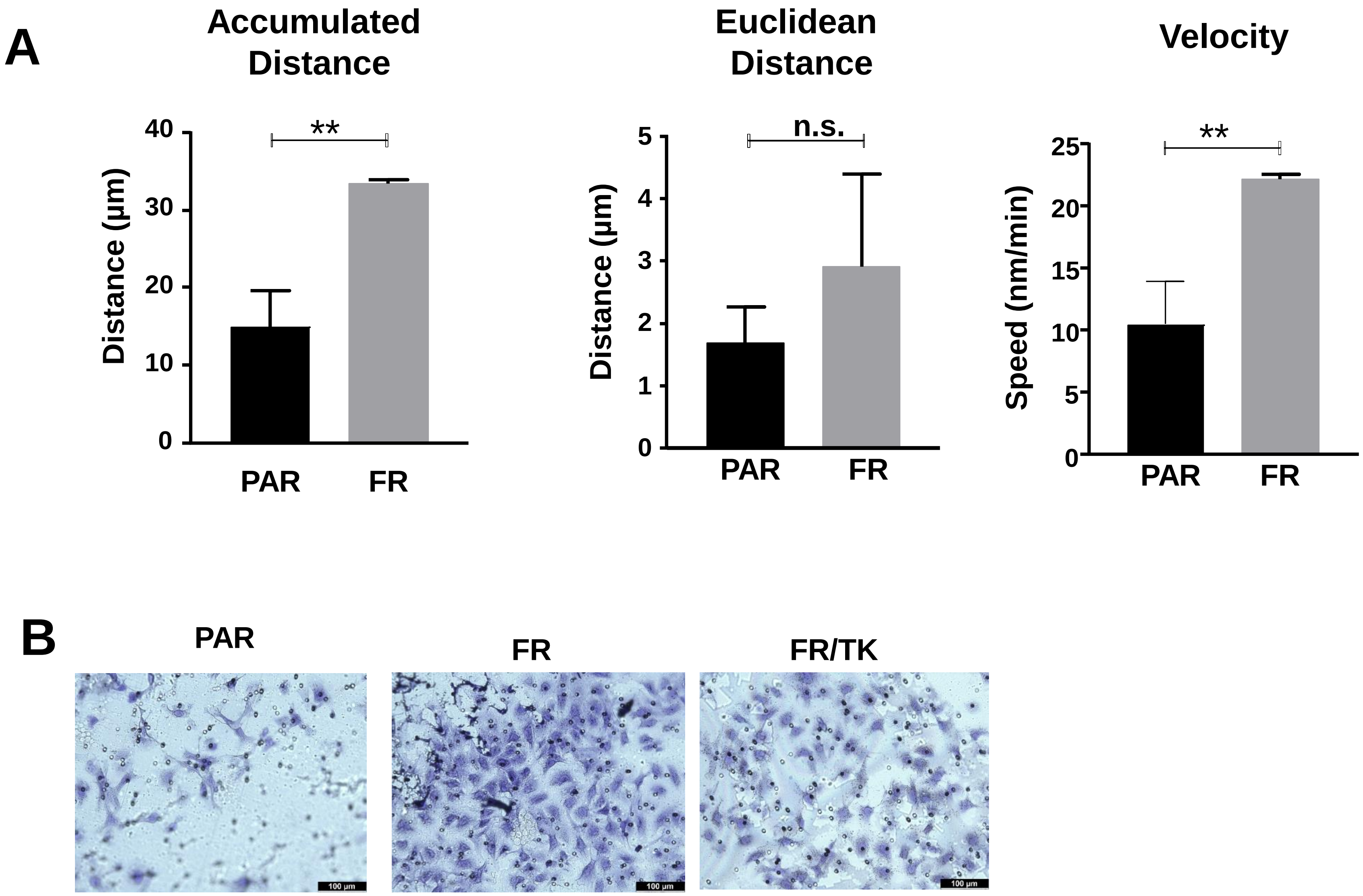


***Supplementary Figure 1:*** A. Cell motility of PAR and FR cells determined by live cell imaging. Cell motility of 9-12 single cells per group was tracked over 24 h. Accumulated (left graph), Euclidean (middle graph) distances as well as cell velocity (right graph) were calculated (n=3, SEM, t-test, n.s.: not significant, ** p<0.01). B. Microphotographs of the lower membrane side of a matrigel invasion chamber 6 h after seeding.
