## Supplementary material for "Development of an optimized, non-stem cell line for intranasal delivery of therapeutic cargo to the central nervous system": Suppl. Fig. 2

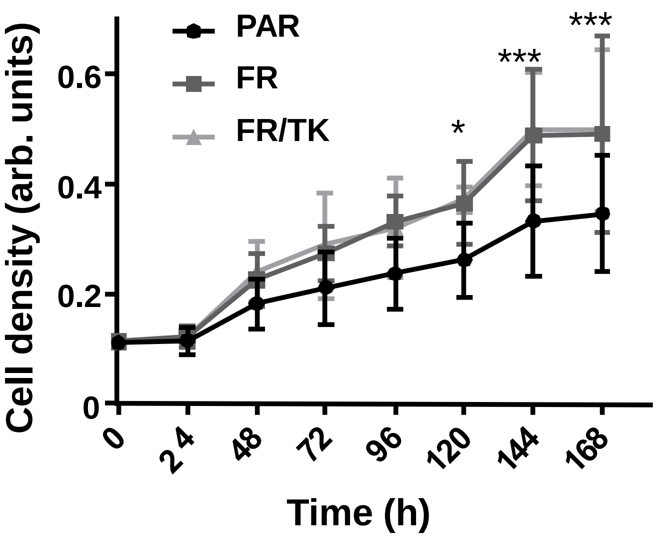


***Supplementary Figure 2:*** Proliferation of PAR, FR and FR/TK cells. 6.000 cells were seeded in microtiter plates and allowed to attach. 4 h after seeding and subsequently every 24 h later cell density was determined by staining the cells with crystal violet (n=3, SEM, t-test, * $p<0.05$; *** $p<0.001$).
