## Supplementary material for "Development of an optimized, non-stem cell line for intranasal delivery of therapeutic cargo to the central nervous system": Suppl. Fig. 3

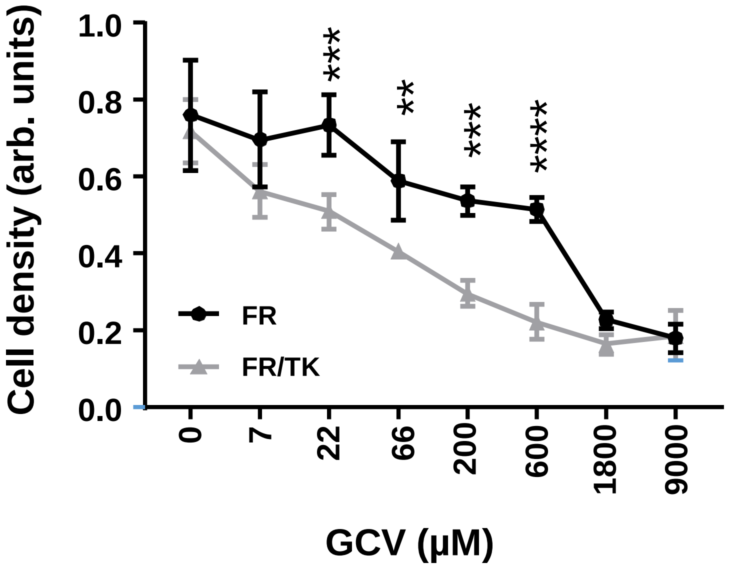


***Supplementary Figure 3:*** GVC vulnerability of FR and FR/TK cells. The cells were seeded in triplicates in microtiter plates and were treated with increasing concentrations of GCV for 72h. Cell density was determined by crystal violet staining (n=3; SEM; 2-way ANOVA, * $p<0.05$, ** $p<0.01$, *** $p<0.001$, **** $p<0.0001$).
