## Supplementary material for "Development of an optimized, non-stem cell line for intranasal delivery of therapeutic cargo to the central nervous system": Suppl. Fig. 4

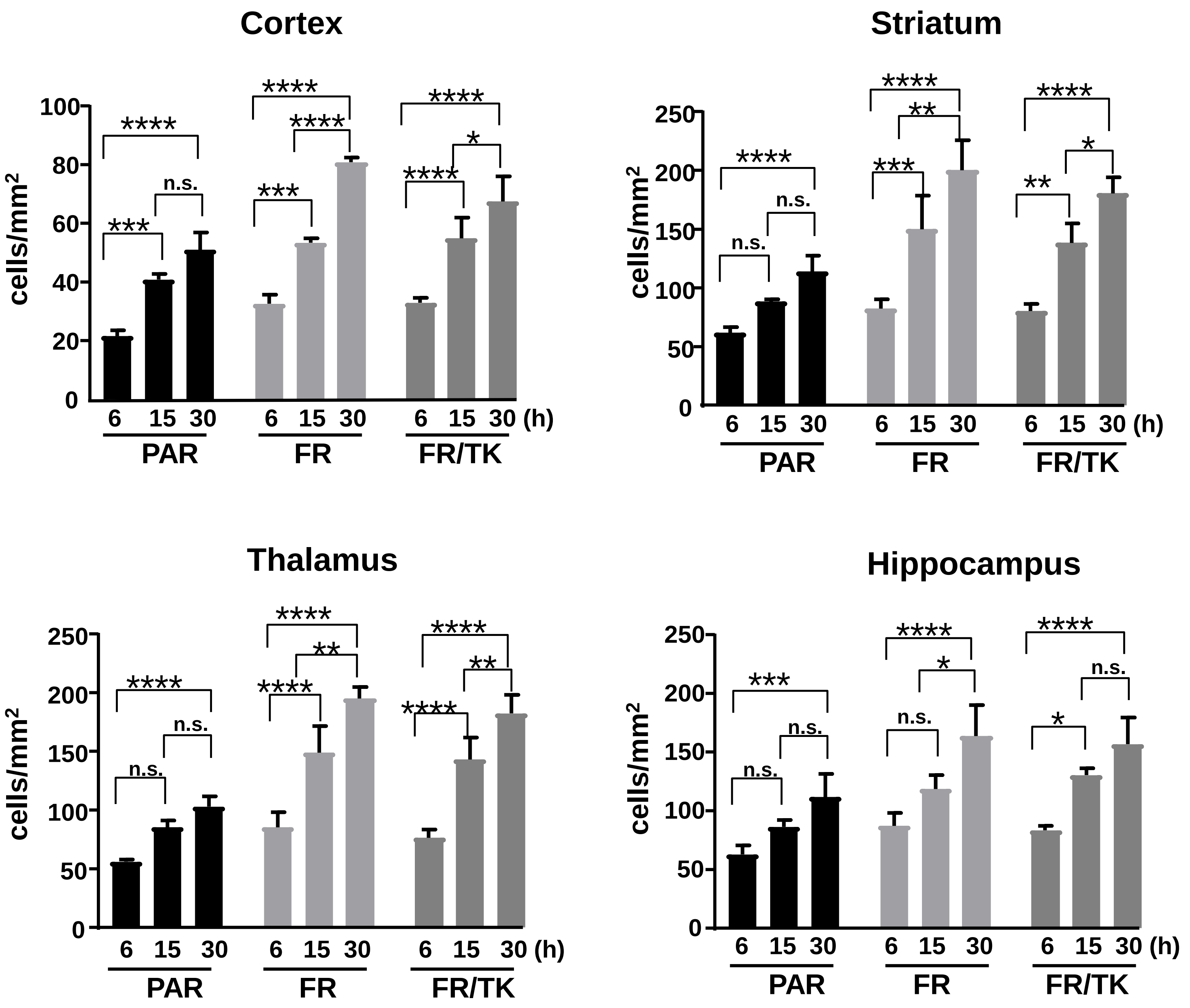


***Supplementary Figure 4:*** NSG mice were treated as indicated in Fig. 3. The enrichment of PAR, FR and FR/TK cells over time in different brain areas is presented in this figure (n=3-4 mice per group, 6-8 slices per mouse and brain area were quantified).
