## Supplementary figures and images for "Development of an optimized, non-stem cell line for intranasal delivery of therapeutic cargo to the central nervous system"

### Suppl. Fig. 5

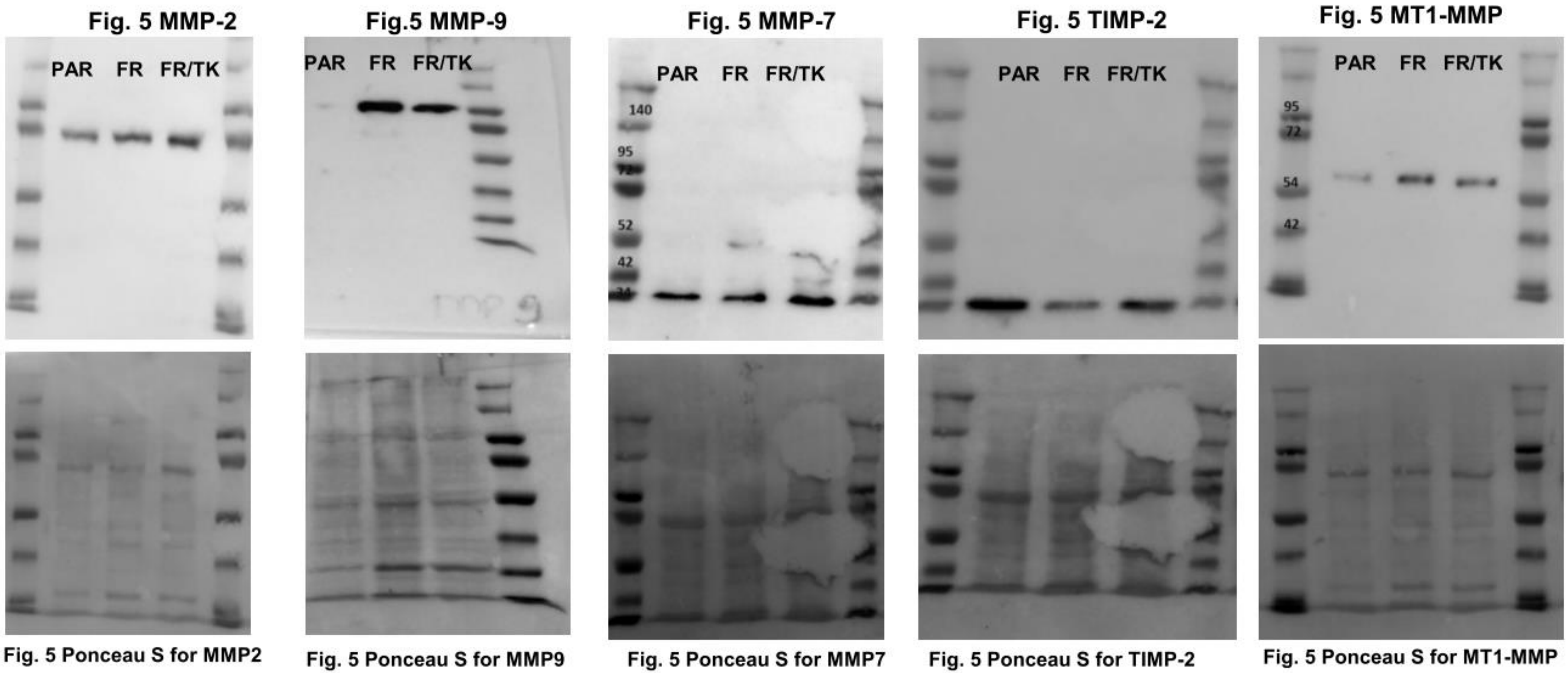


***Supplementary Fig. 5:*** Uncropped immunoblots as shown in partial in Fig. 5
