## Supplementary material for "Development of an optimized, non-stem cell line for intranasal delivery of therapeutic cargo to the central nervous system": Suppl. Fig. 6

Eurofins Genomics Europe Applied Genomics GmbH, Anzinger Str. 7 a, D-85560 Ebersberg

Prof. Dr. Ulrike Naumann
University Tübingen, Hertie Institute
Otfried-Müller-Straße 27
72076 Tübingen
Germany

eurofins | Genomics

SOP_APG_Zelllinienauthentizität_A04_2.0

Person in charge: Dr. Torsten Brendel
Report date: 25.05.2023
Sample received on: 17.05.2023
Start / End of Analysis: 17.05.2023/25.05.2023

### Analytical Report:
### Cell Line Authentication Test
### Order ID: 11108431290

**Method:**

Genetic characteristics were determined by PCR-single-locus-technology.
16 independent PCR-systems D8S1179, D21S11, D7S820, CSF1PO, D3S1358, TH01, D13S317, D16S539, D2S1338, AMEL, D5S818, FGA, D19S433, vWA, TPOX and D18S51 were investigated.
(ASN-0002 core markers are colored grey, Thermo Fisher, AmpFlSTR® Identifiler® Plus PCR Amplification Kit)
In parallel, positive and negative controls were carried out yielding correct results.
Method details are given in **SOP_APG_Zelllinienauthentizität_2.0**

**Result:**

| | **LN-229** | **U87MG** | | | **R49 GSCs** | | **LX-2** | **HB1.F3** | **HEK293** | **HEK293FT** |
|---|---|---|---|---|---|---|---|---|---|---|
| **Client Sample Name** | LN-229 | U87 | [redacted] | [redacted] | R49 GSC pass44 v. 2018 | [redacted] | LX-1 pass 6 | HB1.F3 pass 33 v. 2014 | 293 MX | 293 FT |
| **Sample Code** | CL00013070 | CL00013071 | [redacted] | [redacted] | CL00013068 | [redacted] | CL00013073 | CL00013074 | CL00013075 | CL00013076 |
| D8S1179 | 13,13 | 10,11 | [redacted] | [redacted] | 11,11 | [redacted] | 13,13 | 14,14 | 12,14 | 12,14 |
| D21S11 | 29,30 | 28,32.2 | [redacted] | [redacted] | 29,30 | [redacted] | 28,31 | 28,28 | 28,30.2 | 30.2,30.2 |
| D7S820 | 8,11 | 8,9 | [redacted] | [redacted] | 12,12 | [redacted] | 11,11 | 8,11 | 11,12 | 11,11 |
| CSF1PO | 12,12 | 10,11 | [redacted] | [redacted] | 9,10 | [redacted] | 10,12 | 11,11 | 11,12 | 7,12 |
| D3S1358 | 16,17 | 16,17 | [redacted] | [redacted] | 14,16 | [redacted] | 13,15 | 15,15 | 15,17 | 15,17 |
| TH01 | 9.3,9.3 | 9.3,9.3 | [redacted] | [redacted] | 8,9 | [redacted] | 9.3,9.3 | 6,9 | 7,9.3 | 7,9.3 |
| D13S317 | 10,11 | 11,11 | [redacted] | [redacted] | 11,11 | [redacted] | 11,13 | 8,9 | 12,12 | 12,14 |
| D16S539 | 12,12 | 12,12 | [redacted] | [redacted] | 11,13 | [redacted] | 13,13 | 9,12 | 9,13 | 9,13 |
| D2S1338 | 19,20 | 20,23 | [redacted] | [redacted] | 24,24 | [redacted] | 17,17 | 19,22 | 19,19 | 19,19 |
| D19S433 | 12,17.2 | 15,15.2 | [redacted] | [redacted] | 13,13 | [redacted] | 13,15.2 | 14,14.2 | 15,18 | 15,18 |
| vWA | 16,19,20 | 15,17 | [redacted] | [redacted] | 18,18 | [redacted] | 17,17 | 16,19 | 16,19 | 16,19 |
| TPOX | 8,8 | 8,8 | [redacted] | [redacted] | 8,12 | [redacted] | 8,9 | 10,11 | 11,11 | 11,11 |
| D18S51 | 13,15 | 13,13 | [redacted] | [redacted] | 18,18 | [redacted] | 12,12 | 16,19 | 17,17 | 17,18 |
| AMEL | X,X | X,X | [redacted] | [redacted] | X,X | [redacted] | X,Y | X,X | X,X | X,X |
| D5S818 | 11,12 | 11,12 | [redacted] | [redacted] | 11,11 | [redacted] | 11,12 | 11,12 | 8,9 | 8,9 |
| FGA | 23,23 | 18,24 | [redacted] | [redacted] | 25,25 | [redacted] | 21,26 | 23,24 | 23,23 | 23,23 |
| **Cellosaurus ID:** | **CVCL_0393** | **CVCL_0022** | | | **not available** | | **CVCL_5792** | **CVCL_LJ44** | **CVCL_0045** | **CVCL_6911** |

Supplementary Figure 6: Cell line authentication
